# Impact of Binge Drinking During College on Resting State Functional Connectivity

**DOI:** 10.1101/2021.02.09.430381

**Authors:** Tien T. Tong, Jatin G. Vaidya, John R. Kramer, Samuel Kuperman, Douglas R. Langbehn, Daniel S. O’Leary

## Abstract

**Aim:** The current study aimed to examine the longitudinal effects of standard binge drinking (4+/5+ drinks for females/males in 2 hours) and extreme binge drinking (8+/10+ drinks for females/males in 2 hours) on resting state functional connectivity.

**Method:** 119 college students with distinct alcohol bingeing patterns (35 non-bingeing controls, 44 standard bingers, and 40 extreme bingers) were recruited to ensure variability in bingeing frequency. Resting state fMRI scans were obtained at time 1 when participants were college freshmen and sophomores and again approximately two years later. On four occasions during the 2-year period between scans, participants reported monthly standard and extreme binge drinking for the past 6 months. Association between bingeing and change in functional connectivity was studied using both network-level and edge-level analysis. Network connectivity was calculated by aggregating multiple edges (a functional connection between any two brain regions) affiliated with the same network. The network-level analysis used mixed-effects models to assess the association between standard/extreme binge drinking and change in network connectivity, focusing on canonical networks often implicated in substance misuse. On the other hand, the edge-level analysis tested the relationship between bingeing and change in whole-brain connectivity edges using connectome-based predictive modeling (CPM).

**Results:** For network-level analysis, higher standard bingeing was associated with a decrease in connectivity between Default Mode Network-Ventral Attention Network (DMN-VAN) from time 1 to time 2, controlling for the initial binge groups at time 1, longitudinal network changes, in-scanner motion and other demographic covariates. For edge-level analysis, the CPM failed to identify a generalizable predictive model of cumulative standard/extreme bingeing from change in connectivity edges.

**Conclusions:** Our findings suggest that binge drinking is associated with abnormality in networks implicated in attention allocation and self-focused processes, which, in turn, have been implicated in rumination, craving, and relapse. More extensive alterations in functional connectivity might be observed with heavier or longer binge drinking pattern.

## 1. Introduction

Binge drinking, defined as consuming 4+/5+ drinks for females/males in 2 hours (National Institute on Alcohol Abuse and Alcoholism; NIAAA), is a serious public health issue (Sacks et al., 2015; Stahre et al., 2014). Binge drinking is most common among younger adults (ages 18-34; Kanny et al., 2018), especially those in college (H. R. White et al., 2006). This is problematic, as binge drinking is associated with academic failure, injuries, serious health problems, risky sexual behaviors, memory blackouts, cognitive deficits (Lees et al., 2019; O’Leary et al., 2019; Weschler et al., 1994), and increased risk for developing Alcohol Use Disorder (AUD; Knight et al., 2002).

Researchers have argued that the standard binge drinking definition might be too low to identify heavy episodic drinkers, as both adolescents and young adults commonly drink well beyond this threshold (Hingson and White, 2013; Naimi et al., 2010; Patrick et al., 2013; Read et al., 2008; A. M. White et al., 2006). One suggestion to address this issue is to examine more severe levels of binge drinking, such as extreme binge drinking, which can be defined as consuming 8+/10+ drinks for females/males in 2 hours (Linden-Carmichael et al., 2017; Patrick, 2016; Patrick et al., 2016). Compared to standard bingeing, extreme bingeing is associated with a greater number of negative alcohol-related consequences (impaired control, blackouts, injuries, emergency department visits, legal problems), as well as increased risk for developing AUD (Hingson et al., 2017; Linden-Carmichael et al., 2017; Read et al., 2008).

As standard and extreme binge drinking are important risk factors for AUD development (Knight et al., 2002; Linden-Carmichael et al., 2017), there is significant interest in studying their neurobiological underpinnings. Deficits in various cognitive domains, including verbal memory (Carbia et al., 2018), decision-making (Lees et al., 2019), and inhibitory control (Carbia et al., 2018; Lees et al., 2019) have been linked to binge drinking. However, little is known about abnormal brain connectivity associated with these cognitive deficits. Resting state functional magnetic resonance imaging (fMRI) is a valuable tool for measuring intrinsic functional connectivity by assessing the correlation of the blood-oxygen-level-dependent (BOLD) signal fluctuations in different brain regions (Biswal et al., 1995). Resting state connectivity networks are replicable (Power et al., 2011; Seitzman et al., 2020; Yeo et al., 2011) and have the potential to be used as biomarkers for substance abuse and dependence (Zilverstand et al., 2018).

Most studies on the links between binge drinking and resting state functional connectivity have focused on the reward, salience, and executive control networks. The most replicated finding is the negative association between alcohol misuse and connectivity within the reward network, specifically, between amygdala and orbitofrontal cortex (OFC). Lower amygdala-OFC connectivity was associated with higher alcohol use in both cross-sectional (Peters et al., 2015) and longitudinal analyses (Peters et al., 2017). In addition, lower amygdala-OFC connectivity was associated with higher weekly drinking in the past month (Crane et al., 2018). On the other hand, when assessing different seeds of the reward network (i.e., connectivity between nucleus accumbens (NAcc)-OFC, instead of amygdala-OFC), an opposite pattern was reported such that compared to young adult light drinkers, binge drinkers showed increased NAcc-OFC connectivity (Arienzo et al., 2019).

Functional connectivity between the salience and other networks has also been associated with alcohol misuse. Arienzo and colleagues (2019) have reported increased connectivity between the salience and the habit network (dorsal anterior cingulate cortex (dACC)-caudate) in binge drinkers relative to light drinkers. In addition, salience-reward (dACC-amygdala) connectivity was shown to negatively correlate with alcohol misuse in a group of non-dependent alcohol drinkers (Hu et al., 2018). Another commonly studied resting state network in binge drinking is the executive control network. However, findings on the direction of the association between alcohol misuse and executive control network connectivity are inconsistent. Sousa and colleagues (2019) reported increased connectivity within the executive control network in bingers, as compared to alcohol abstinent controls. On the other hand, decreased connectivity within the executive control network was found when comparing 87 controls with 383 problematic alcohol users whose drinking patterns ranged from binge drinking to severe AUD (Weiland et al., 2014). Lastly, decreased connectivity between executive control and memory network (inferior frontal gyrus-hippocampus) was also reported in binge drinkers (Arienzo et al., 2019).

Abnormal connectivity of networks other than the reward, salience, and executive control networks have also been reported in binge drinkers. A longitudinal resting state magnetoencephalography (MEG) study found that compared to controls, college binge drinkers who maintained a binge drinking pattern for two years showed an increased Default Mode Network connectivity (DMN; Correas et al., 2016). Lastly, greater binge drinking was significantly correlated with decreased Ventral Attention Network connectivity (VAN) in a group of college students with varying levels of binge drinking (Herman et al., 2019).

Previous studies in the literature have several shortcomings. Firstly, most studies were cross-sectional (except Correas et al., 2016; Peters et al., 2017), making it difficult to tease apart predisposing factors vs. potential consequences of binge drinking. Secondly, most studies used a seed-based method to assess resting state functional connectivity, which has many limitations. In particular, the seed-based method limits research replicability, as most researchers choose to focus on unique seeds and often do not attempt to replicate earlier results. This can result in inconsistent findings such as the positive (Arienzo et al., 2019) and negative (Crane et al., 2018; Peters et al., 2017, 2015) associations between alcohol misuse and connectivity within the reward network. This apparent inconsistency may have stemmed from using different seeds (NAcc, OFC, amygdala) affiliated with the reward network. Lastly, results from the seed-based method are often limited to canonical networks frequently implicated in addiction. This could overlook important information from the whole-brain functional connectome. When whole-brain connectivity is considered, aberrant connectivity in non-canonical networks in addiction such as visual or sensorimotor networks have emerged (Fede et al., 2019; Ruan et al., 2019).

To address these shortcomings, the current study longitudinally assessed change in resting state functional connectivity as a function of standard and extreme binge drinking frequency reported during the 2-year period after the first MRI session. In addition, instead of choosing a small number of seeds as in the seed-based approach, we used the Seitzman atlas (Seitzman et al., 2020), which included 300 seeds affiliated with 13 networks. This atlas is the extension to the Power atlas (Power et al., 2011), which only included cortical seeds, so that connectivity of subcortical and cerebellar seeds were also examined. We hypothesized that the relationship between binge drinking and functional connectivity will mirror connectivity abnormalities found in addiction, as outlined by Zilverstand and colleagues (Zilverstand et al., 2018). Therefore, we focused on the following eights networks often implicated in addiction: Cingulo-Opercular (CO), Default Mode Network (DMN), Dorsal Attention Network (DAN), Fronto-Parietal (FP), Medial Temporal Lobe (MTL), Reward, Salience, and Ventral Attention Network (VAN). In the network-level analysis, mixed-effects models were used to test the association between alcohol bingeing and network connectivity, which was calculated by aggregating multiple edges of the same network (an edge is a connectivity between two seeds). This network-level analysis allowed us to 1) test the association between bingeing and connectivity in a theory-driven fashion and 2) assess network connectivity more objectively, as the network connectivity calculation took into account all seeds of that network. The latter point addressed one limitation of the seed-based method, in which researchers often picked one or two seeds to represent the connectivity of the whole network. The network-level analysis does have some limitations. Firstly, as mentioned previously, focusing solely on canonical networks can lead to overlooking non-canonical connectivity that might also be important in understanding binge drinking. Secondly, although the network connectivity calculation is more objective than picking a small number of seeds to represent a network, simply averaging across all edges of a network implies that these edges contribute equally to network connectivity, which is not necessarily accurate. Therefore, we also used the connectome-based predictive modeling (CPM; Shen et al., 2017), which is a data-driven edge-level analysis that can address both issues. That is, CPM extends the analysis to non-canonical networks and only focuses on important edges that significantly associated with binge drinking. Lastly, to better understand the behavioral correlates of connectivity variables, correlations between change in cognitive performance and change in connectivity were calculated for connectivity variables that were significantly associated with standard/extreme bingeing.

## 2. Material and Methods

### 2.1. Participants

The current study is a part of a larger project on binge drinking and brain maturation (O’Leary et al., 2019; Tong et al., 2020). To ensure variability in levels of binge drinking, we recruited 3 groups of participants that were all right-handed and had distinct baseline bingeing patterns: non-bingeing controls (Control), standard bingers (sBinge), and extreme bingers (eBinge). Group membership was based on a screening questionnaire developed by our lab, which participants completed prior to the first in-lab session. To be included in the Control group, participants had to have no history of cannabis use or standard/extreme bingeing, but non-binge alcohol consumption (1-2 drinks/occasion) was allowed. To be included in the sBinge/eBinge groups, participants must have had at least 2 sBinge/eBinge episodes, respectively, in the past 30 days or since the semester began. Additionally, to minimize potential effects of marijuana, sBinge/eBinge participants must have had no more than 3 occasions of marijuana use in the past month, and no more than 30 lifetime uses. Both bingeing groups also had to have limited lifetime use (<15 times) of other substances except nicotine. Other exclusion criteria included: history of seizure disorders, head injury, neurologic, metabolic, or cardiovascular disease, cerebrovascular events, or meeting DSM-IV criteria for major psychiatric disorders (including substance use disorders other than alcohol use disorder for both binge groups), based on the Mini International Neuropsychiatric Interview (Hergueta et al., 1998). In the two MRI sessions, all subjects passed a breathalyzer test and had negative urine screens for all drugs except for MJ (due to its long half-life). Out of the 161 participants that completed the first session, 119 participants (35 Control, 44 sBinge, and 40 eBinge) came back for the second session. No one met the exclusionary criterion for excessive in-scanner movements (having at least one frame-to-frame movement greater than 3mm in any direction).

### 2.2. Procedures

All procedures were approved by the University of Iowa Institutional Review Board. The procedures were the same in the two in-lab sessions: All participants signed informed consent forms, completed a brief medical history screening, web-based self-report measures, and underwent the structural and resting state fMRI scans. The two sessions were approximately two years apart. During the 2-year gap, participants were asked to complete an online survey every six months. The online surveys assessed monthly binge and extreme binge frequency. For example, six months after the first session, participants were asked to report binge and extreme binge frequency (days/month) for each of the prior six months.

### 2.3. FMRI Acquisition

All MRI data at time 1 were collected with a Siemens 3T scanner. At time 2, data from 21 participants were collected using the same Siemens scanner, and data from 98 participants were collected using a 3T General Electric (GE) scanner. For the Siemens scanner, the following parameters were used: T1-weighted (TR=2,530 ms, TE=2.8 ms, flip angle=10 degrees, voxel size=1.0×1.0×1.0 mm, frames=256, FOV=256×256), resting state (TR=2,000 ms, TE=30 ms, flip angle=77 degrees, voxel size=3.4×3.4×3.5 mm, frames=180, FOV=384×384, scan duration=360 s). For the GE scanner, the following parameters were used: T1-weighted (TR=8.46 ms, TE=3.248 ms, flip angle=12 degrees, voxel size=1.0×1.0×1.0 mm, frames=252, FOV=256×256), resting state (TR=2,000 ms, TE=30 ms, flip angle=77 degrees, voxel size=3.4×3.4×3.5 mm, frames=5580, FOV= 64×64, scan duration=360 s). Scanner type was added as a covariate in all analyses.

### 2.4. Neuropsychological Tests

Four tasks from the Computerized Multiphasic Interactive Neurocognitive Dual Display System (CMINDS) were administered: Stroop (executive function – interference), Digit Span (attention), Letter Fluency (language), and Trail Making (Trail A: attention, Trail B: executive function – sequencing). The Stroop task consisted of 4 conditions: 1) read out loud a list of color names printed in black ink, 2) name out loud the colors of a set of colored squares, 3) say out loud the color that a set of non-color words are printed in, and 4) say out loud the color that a set of words are printed in, but the ink color is incongruent with the color word. On the Digit Span task, participants viewed digit sequences and reported from memory these sequences either in the same (Forward) or reverse (Backward) order. The task started with a sequence of two digits, and new digits would be added after two successful trials at a given length. If participants failed both sequences at a given length, or if the maximum span (Forward: 9 digits, Backward: 8 digits) was reached, the task would end. In the Letter Fluency task, participants were asked to say in 1 minute as many words as they could that start with a specified letter. Lastly, in the Trail Making task, participants were presented with a two-dimensional graphic that comprised of either numbered circles (Trail A), or of circled numbers and letters (Trail B). They were instructed to draw lines connecting the circles of either consecutive numbers (Trail A) or alternating between consecutive numbers and consecutive letters (Trail B; e.g., 1-A, 2-B, etc.). The variables of interest were Stroop Effect 1 (naming ink color of incongruent color words minus naming colors of colored squares), Stroop Effect 2 (naming ink color of incongruent color words minus naming ink color of non-color words), digit span total correct (forward + backward), letter fluency total correct, and trail making elapsed time (Trail B minus Trail A).

### 2.2. Statistical Analysis

#### 2.5.1. Group Comparisons

Since participants were recruited in three baseline groups to ensure variability in binge/extreme binge drinking, group comparisons were conducted to characterize the participants. Group differences in sex, race, and parental socioeconomic status (SES) were assessed with chi-square tests. Cumulative standard and extreme bingeing were calculated by adding up monthly bingeing reported during the 24 months after session 1. Group differences in cumulative standard and extreme bingeing were tested using Kruskal-Wallis tests and followed up with Dunn pair-wise tests with False Discovery Rate (FDR) correction for multiple comparisons. Lastly, group differences in age and framewise displacement (FD) were tested for each session separately using ANOVA.

#### 2.5.2. FMRI Preprocessing

Prior to all analysis, the first three volumes were discarded to remove artifacts associated with scanner disequilibrium. Afterward, all preprocessing steps were run with fMRIPrep version 1.4.1 (Esteban et al., 2019). The anatomical workflow included the following steps: 1) correcting for intensity non-uniformity (Tustison et al., 2010) of the 2 T1w images with ANTs 2.2.0 (Avants et al., 2009); 2) creating a reference T1w by registering these two INU-corrected T1w images using FreeSurfer 6.0.1 (Reuter et al., 2010), segmenting the reference T1w using FAST (Zhang et al., 2001); and 3) non-linearly normalizing the reference T1w to a standard space (FSL’s MNI ICBM 152 non-linear 6th Generation Asymmetric Average Brain Stereotaxic Registration Model) using ANTs 2.2.0 (Evans et al., 2012). For the functional images, the following steps were applied: 1) motion correction using MCFLIRT (FSL 5.0.9; Jenkinson et al., 2002); 2) slice time correction using 3dTshift from AFNI 20160207 (Cox and Hyde, 1997); 3) spatial smoothing with a Gaussian kernel of 6mm full-width half-maximum, co-registering to MNI; and 4) non-aggressive denoising with ICA-AROMA (Pruim et al., 2015).

After preprocessing with fMRIPrep, nuisance regression and bandpass filtering (0.009-0.08 Hz) were done with AFNI 3dTproject. Nuisance regressors included mean signal from the white matter and cerebrospinal fluid masks (95% probability masks) extracted from the non-aggressively denoised, unsmoothed BOLD. Lastly, the data were normalized and scaled.

#### 2.5.3. Network-Level Connectivity

AFNI 3dNetCorr (Taylor and Saad, 2013) was used to construct the Fisher Z-transformed 300×300 correlation matrix for each participant. Within-network connectivity variables were calculated by averaging connectivity strengths of all edges that affiliated with the same networks. Between-network connectivity variables were calculated by averaging the correlations of all seed-to-seed connections between different networks. By default, 3dNetCorr did not calculate the correlation of any seed with more than 10% of voxels with null times series. In addition, seeds with few data points might have high between-subject variance, causing instability in the within- and between-network connectivity calculation. Therefore, 15 seeds with high signal dropout (see Inline Supplementary Table 1) were excluded for this analysis. However, because of their hypothetical importance in addiction, we excluded the following regions from the aforementioned signal dropout rules: amygdala, NAcc, and hippocampus. The seed with the fewest data points (213/238 runs) included in this analysis was the right amygdala.

From the 8 *a priori* chosen networks (CO, DMN, DAN, FP, MTL, Reward, Salience, and VAN), 8 within-network variables and 28 between-network variables were the outcome variables of the mixed-effects models testing the association between binge/extreme binge drinking and change in network connectivity. For each of these outcome variables, the fixed effects of interest included cumulative standard and extreme bingeing (log-transformed) and group (Control, sBinge, eBinge). We added additional fixed effects to control for potential confounding by baseline age, session (time 1 versus time 2), baseline age × session interaction, sex, scanner type (Siemens, GE), SES, and FD. Participant identifiers were added as random intercepts. Since the log-transformed cumulative standard and extreme bingeing were highly correlated (*r*=.80), these two variables were not entered in the same model. FDR correction for multiple comparison was run separately for each fixed effect of interest (cumulative standard/extreme bingeing and group).

#### 2.5.4. Edge-Level Connectivity

The connectome-based predictive modeling (CPM) was used to examine the association between standard and extreme bingeing with each unique element from the 300×300 correlation matrix (i.e., an edge representing the connectivity between any seed pair). The protocol was based on previous published methodology (Shen et al., 2017) and was run separately for standard and extreme bingeing. Connectivity changes were first calculated by subtracting time 1 from time 2. Input data were thus composed of change in connectivity edges, cumulative standard/extreme bingeing, and confound variables including sex, SES, group, scanner (same or different scanners in the two sessions), and change (time 2 minus time 1) in age and FD. These input data were split into training/test sets using leave-one-out cross validation. The CPM protocol involved the following steps: 1) select important edges whose change in connectivity strength were significantly correlated with standard/extreme bingeing in the training set using Spearman partial correlation, 2) fit a linear model using the training set and apply this model to predict standard/extreme bingeing frequency from change in connectivity edges in the test set, 3) calculate model accuracy using the test sets across all cross-validation folds, and 4) determine if this model accuracy is significant using 500 permutations of the data to create the null distribution.

For Step 1 (feature selection), edges that were positively or negative correlated with standard/extreme bingeing (i.e., positive and negative edges) were analyzed separately and used in separate models to predict bingeing. For Step 2 (model fitting), standard/extreme bingeing were the outcome variables, and for each subject, the edge strengths (z-transformed correlation coefficients) were summed across the important edges identified from Step 1 to calculate the predictor of the model. Model accuracy (Step 3) was the correlation coefficient between the predicted and observed bingeing in the test set across all cross-validation folds (Rpos: coefficient for the model with positive edges, Rneg: coefficient for the model with negative edges). Since the CPM protocol didn’t specify which p-value threshold should be used for Step 1, several thresholds (.01, .001, .0001) were explored. As suggested by Shen and colleagues (2017), *p*<.0001 was chosen, as models using this threshold showed the highest accuracy in predicting bingeing in the training set across all cross-validation folds (see Inline Supplementary Table 2).

While the whole CPM protocol allowed us to test edges whose change in connectivity strength can predict prospective bingeing frequency, Step 1 from CPM allowed us to examine consistent patterns of the association between edge connectivity changes and bingeing. In other words, the goal of the full CPM protocol was to test predictive models, whereas exploring results from Step 1 of CPM provided a detailed description of edge-bingeing associations. Specifically, the leave-one-out cross-validation created small perturbations of the sample, as there were 119 folds of cross-validation, and in each fold, 1 participant was excluded from the training set. Thus, the excluded participant was not used to calculate the correlation between bingeing and edge connectivity changes. Edge connectivity changes were defined as showing a consistent pattern of association with cumulative bingeing if they strongly correlated with bingeing (*p*_uncorrected_<.0001) in all 119 cross-validation folds.

## 3. Results

### 3.1. Group Comparison

The group comparison results are reported in Table 1. Chi-square tests revealed no significant group differences in sex, race, or parent SES. There were also no significant group differences in age at time 1 *(F(2,* 116)=2.32, *p*=0.10), FD at time 1 (*F*(2, 116)=2.41, *p*=0.09) or time 2 (*F*(2, 116)=2.32, *p*=0.10). Lastly, even though groups were classified based on bingeing patterns prior to session 1, there were significant group prospective differences on both cumulative standard bingeing (*χ*^2^(2)=53.53, *p*<.001; see Inline Supplementary Figure 1A) and extreme bingeing (*χ*^2^(2)=54.51, *p*<.001; see Inline Supplementary Figure 1B). Both binge groups reported higher cumulative standard bingeing than the Control group (*p*s<.001), but they did not differ from one another (*p*=.18). On the other hand, for cumulative extreme bingeing, Controls reported lower frequency than sBinge (*p*<.001), who in turn reported lower frequency than eBinge (*p*=.006).

**Table 1.**
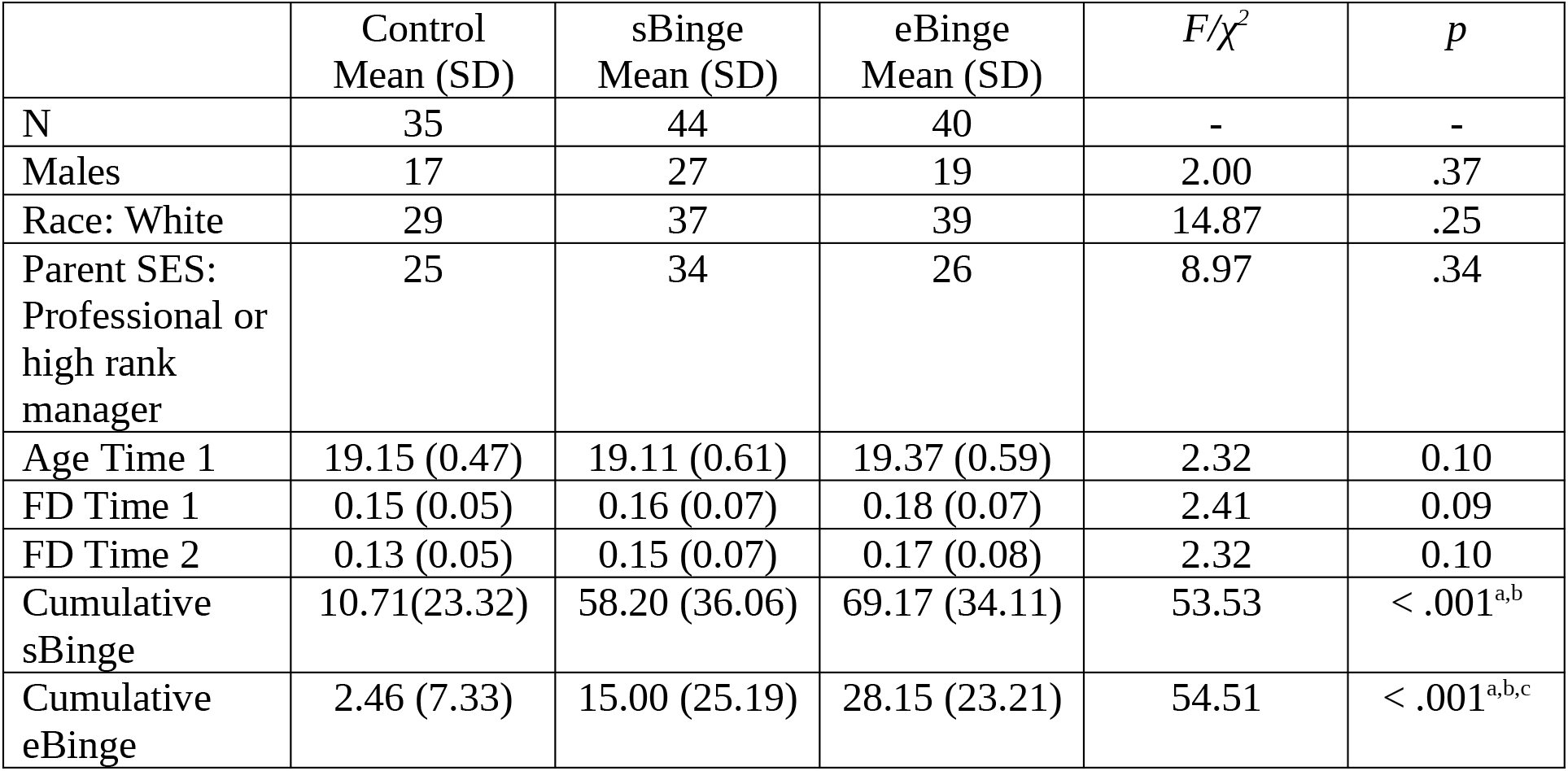
Participant Characteristics. Group differences on Sex, Race, Socioeconomic Status (SES) were assessed with chi-square tests. Group differences on Age and Framewise Displacement (FD) were assessed separately for time 1 and time 2 with ANOVA. Lastly, group differences on cumulative standard bingeing (sBinge) and cumulative extreme bingeing (eBinge) were assessed with Kruskal-Wallis and followed up with pair-wise Dunn test with False Discovery Rate adjustment for multiple comparison. Significant pairwise group test notations: a) Control < sBinge, b) Control < eBinge, and c) sBinge < eBinge.

| | Control<br>Mean (SD) | sBinge<br>Mean (SD) | eBinge<br>Mean (SD) | $F/\chi^2$ | $p$ |
| --- | --- | --- | --- | --- | --- |
| N | 35 | 44 | 40 | - | - |
| Males | 17 | 27 | 19 | 2.00 | .37 |
| Race: White | 29 | 37 | 39 | 14.87 | .25 |
| Parent SES:<br>Professional or<br>high rank<br>manager | 25 | 34 | 26 | 8.97 | .34 |
| Age Time 1 | 19.15 (0.47) | 19.11 (0.61) | 19.37 (0.59) | 2.32 | 0.10 |
| FD Time 1 | 0.15 (0.05) | 0.16 (0.07) | 0.18 (0.07) | 2.41 | 0.09 |
| FD Time 2 | 0.13 (0.05) | 0.15 (0.07) | 0.17 (0.08) | 2.32 | 0.10 |
| Cumulative<br>sBinge | 10.71(23.32) | 58.20 (36.06) | 69.17 (34.11) | 53.53 | < .001<sup>a,b</sup> |
| Cumulative<br>eBinge | 2.46 (7.33) | 15.00 (25.19) | 28.15 (23.21) | 54.51 | < .001<sup>a,b,c</sup> |

### 3.2. Network-Level Connectivity

Significant fixed effect (uncorrected) of cumulative standard bingeing on network connectivity from the 36 models are shown in Table 2.

**Table 2.** Fixed effects of cumulative standard bingeing (sBinge) on network connectivity. FDR: False Discovery Rate. Network abbreviations: Cingulo-Opercular (CO), Default Mode Network (DMN), Salience (Sal), and Ventral Attention Network (VAN).

| | $\beta$ | SE | $t$ | DF | Uncorrected $p$ | FDR |
| --- | --- | --- | --- | --- | --- | --- |
| **CO-VAN** | 0.01 | 0.004 | 2.11 | 163.39 | 0.04 | 0.49 |
| **DMN-VAN** | -0.01 | 0.004 | -3.45 | 168.11 | <0.001 | 0.03 |
| **Sal-VAN** | 0.01 | 0.004 | 1.99 | 156.28 | 0.049 | 0.49 |

Only the fixed effect of cumulative standard bingeing on DMN-VAN connectivity survived the FDR correction *(β* = −0.01, *t*(168.11)= −3.45, *p*_uncorrected_<.001, FDR=.03). In particular, controlling for the effects of group, session, and other covariates, higher standard bingeing was associated with a negative change in DMN-VAN connectivity (Time 2 < Time 1; Figure 1). That is, at lower standard bingeing levels, DMN-VAN connectivity at Time 2 is greater than Time 1. The inverse pattern is true at higher standard bingeing levels.

**Figure 1.**
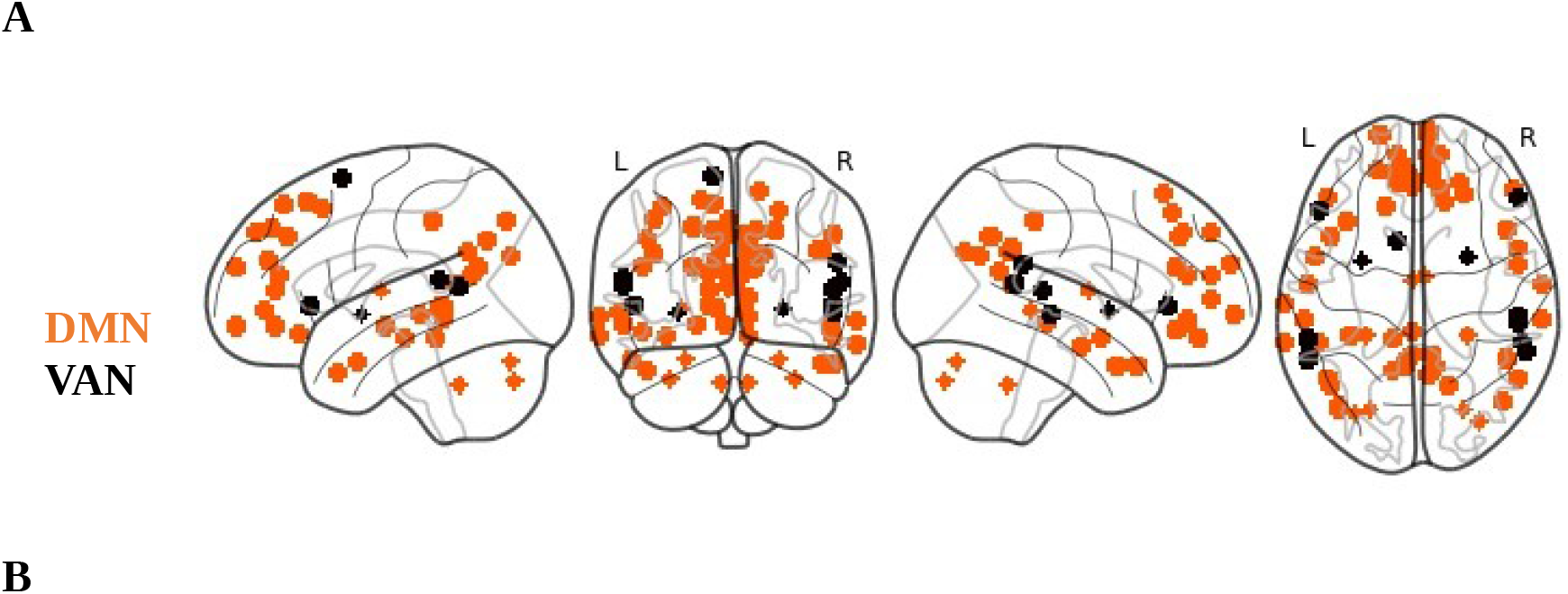
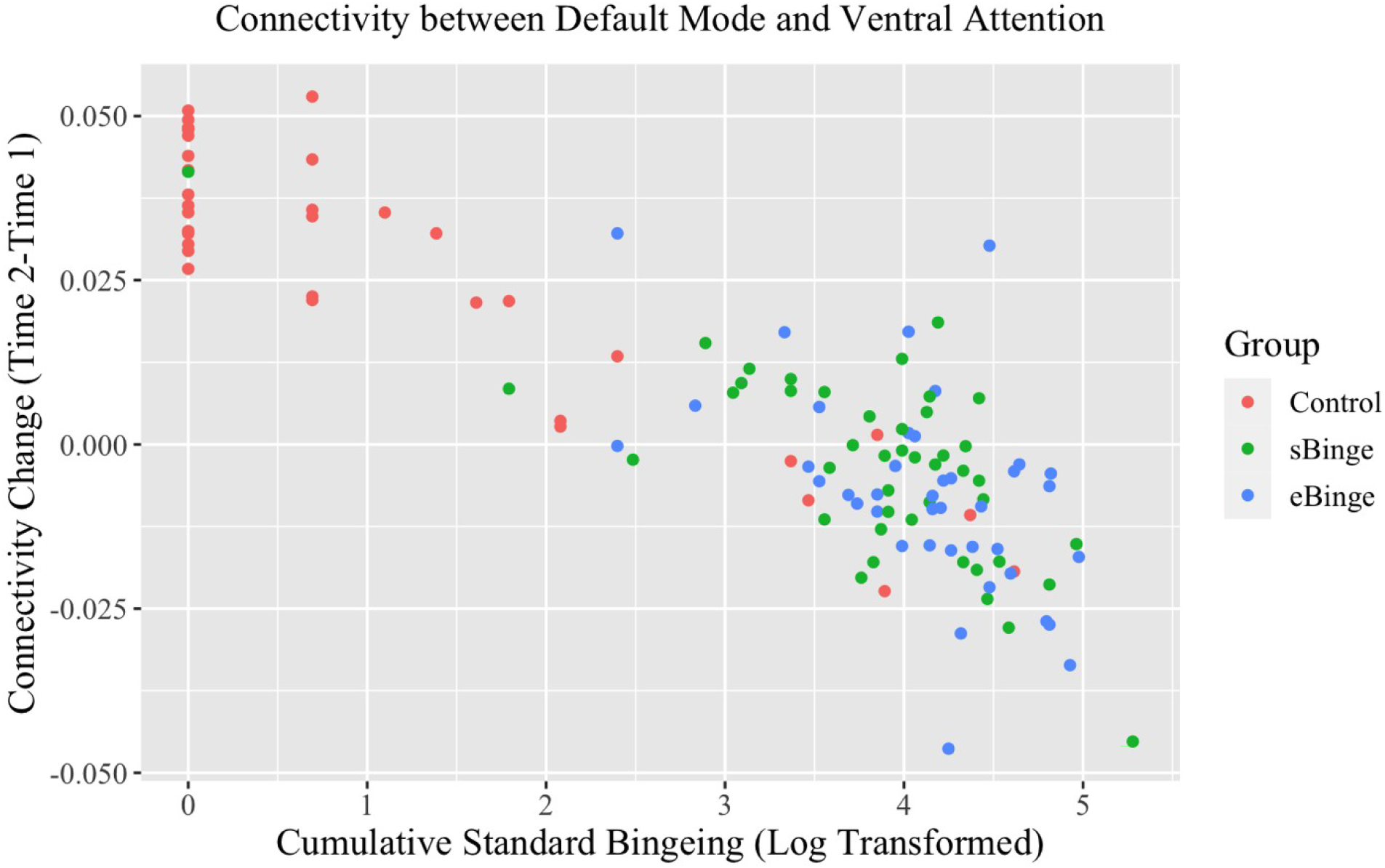
Results of the network-level analysis. **(A)** Seeds of the Default Mode Network (DMN, orange) and Ventral Attention Network (VAN, black). **(B)** Association between predicted change in DMN-VAN connectivity (Time 2 minus Time 1) and log-transformed cumulative standard bingeing.

Significant fixed effect (uncorrected) of cumulative extreme bingeing on network connectivity from the 36 models are shown in Table 3. None of the effects of cumulative extreme bingeing on network connectivity survived the FDR correction. Additionally, fixed effects of cumulative bingeing (summation of standard and extreme bingeing) on the 36 network connectivity variables were also explored, but none of the associations survived the FDR correction (see Inline Supplementary Table 3).

**Table 3.**
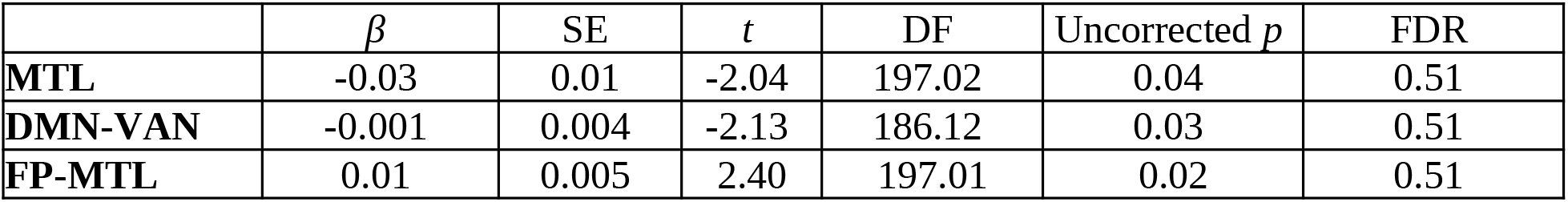
Fixed effects of cumulative extreme bingeing (eBinge) on network connectivity. Network abbreviations: Medial Temporal Lobe (MTL), Default Mode Network (DMN), Ventral Attention Network (VAN), and Fronto-Parietal (FP).

| | $\beta$ | SE | $t$ | DF | Uncorrected $p$ | FDR |
| --- | --- | --- | --- | --- | --- | --- |
| **MTL** | -0.03 | 0.01 | -2.04 | 197.02 | 0.04 | 0.51 |
| **DMN-VAN** | -0.001 | 0.004 | -2.13 | 186.12 | 0.03 | 0.51 |
| **FP-MTL** | 0.01 | 0.005 | 2.40 | 197.01 | 0.02 | 0.51 |

Pearson correlations were computed for the change (Time 2 minus Time 1) in DMN-VAN connectivity and change in cognitive performance (Table 4). Across all 4 cognitive tasks, change in DMN-VAN connectivity was not significantly correlated with change in performance (all *p*_uncorrected_ > .22). Spearman’s correlation between change in cognitive performance and cumulative standard/extreme bingeing were also explored, but none were significant (all *p*_uncorrected_ > .12, see Inline Supplementary Table 4).

**Table 4.**
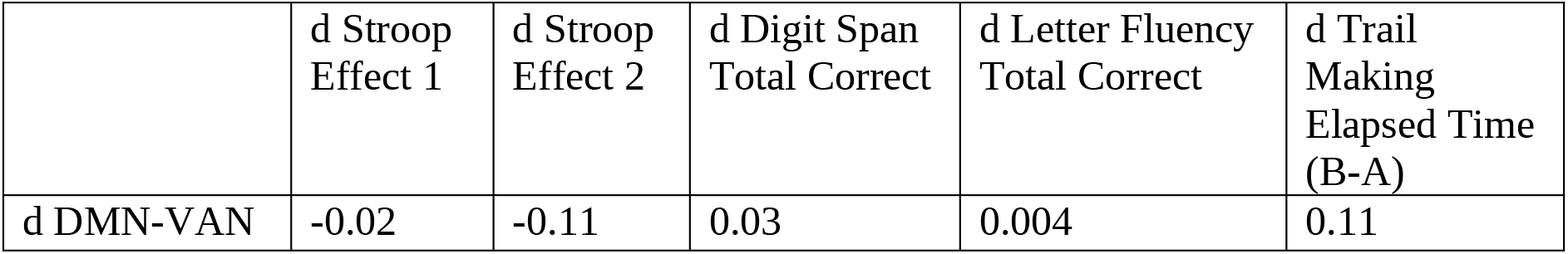
Pearson’s correlation coefficients between change (d = Time 2 minus Time 1) in cognitive performance and change in connectivity between Default Mode Network (DMN) and Ventral Attention Network (VAN). No correlation was significant.

### 3.3. Edge-Level Connectivity

Before testing if change in connectivity edges can predict standard/extreme bingeing using CPM, consistent patterns of the associations between bingeing and edge connectivity changes (or consistency analysis for short) were first explored. As mentioned previously, important edge connectivity-bingeing associations may be lost in the network-level analysis, as all edges contributed equally to the network connectivity calculation. The consistency analysis, despite being exploratory, can be valuable as it highlights important edges that demonstrate significant (*p*_uncorrected_<.0001) and consistent associations with bingeing (see Inline Supplementary Figure 2). The adjacency matrices in Supplementary Figure 2 suggest that the association patterns between edge connectivity changes and standard/extreme bingeing were highly similar, which is not surprising, given the high correlation between standard and extreme bingeing. One way to better understand these results is to focus on important seeds and their associated edges. Important seeds were defined as seeds with high degree calculated from the adjacency matrices. A seed’s degree is the total number of edges linked to that seed. A range of degree threshold was tested, and this section reported results from using the degree threshold of 30 due to its high interpretability. Specifically, patterns of consistent positive associations between edge connectivity changes and standard (Figure 2A) and extreme bingeing (Figure 2C) were highly similar, with important seeds affiliated with the DMN and the Somatomotor Dorsal Network. Interestingly, the consistency analysis between standard bingeing and edge connectivity changes revealed important seeds in the DMN and the VAN, which is somewhat consistent with the network-level analysis. However, this analysis also revealed an important seed in the Somatomotor Dorsal Network, a non-canonical addiction network, emphasizing the importance of studying the whole-brain connectome. On the other hand, the patterns of consistent negative associations between edge connectivity changes and standard/extreme bingeing were less consistent. Important seeds for negative associations between edge connectivity changes and standard bingeing included seeds in the Auditory, CO, DAN, Salience, and Somatomotor Dorsal Network (Figure 2B). Important seeds for negative associations between edge connectivity changes and extreme bingeing included seeds in the CO and DMN (Figure 2D).

**Figure 2.**
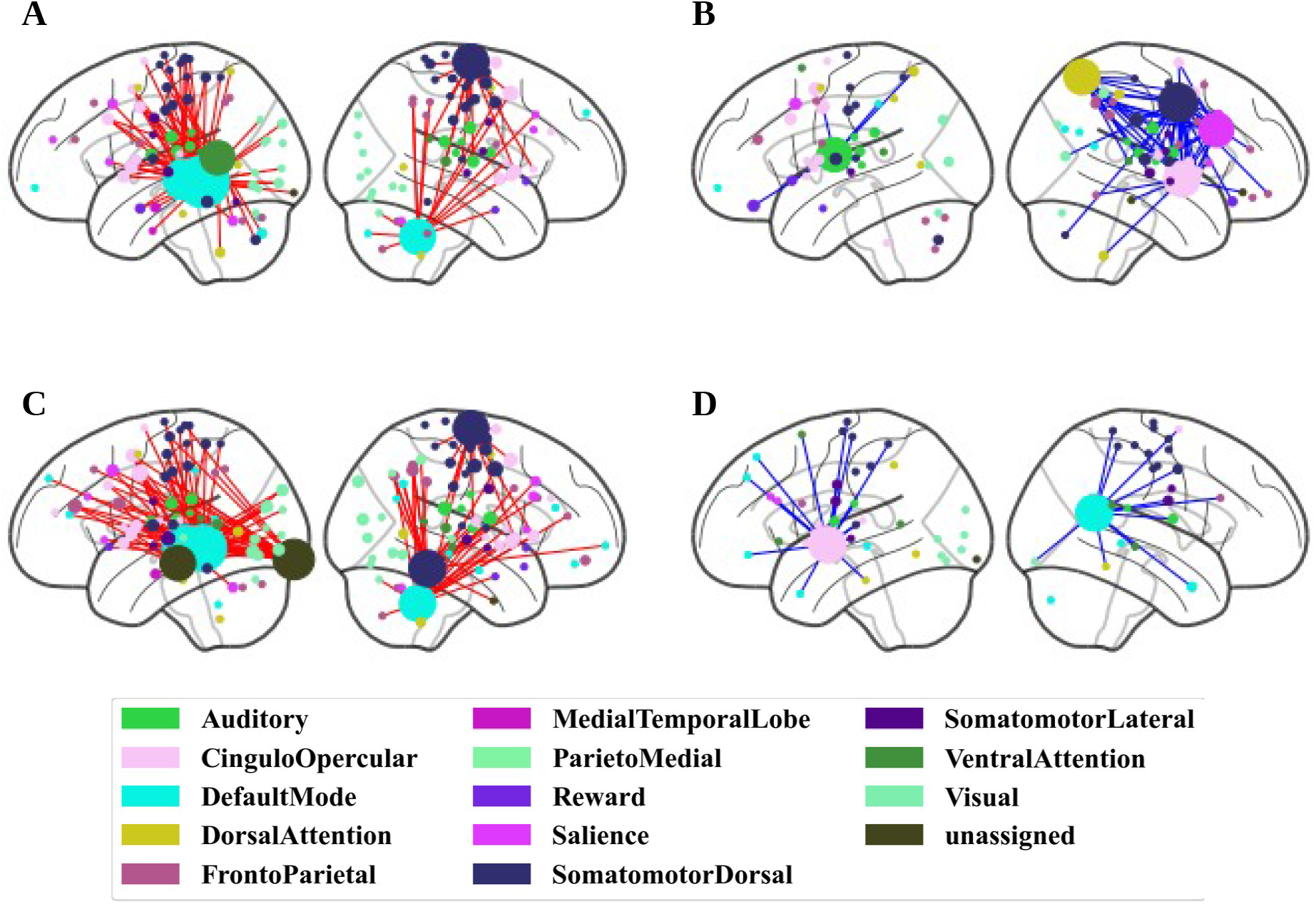
Results of the consistency analysis. Edges with change in connectivity strength (Time 2 minus Time 1) that demonstrated significant (*p*_uncorrected_<.0001) and consistent associations with cumulative standard bingeing (A) positive and B) negative associations) and cumulative extreme bingeing (C) positive and D) negative associations). Only important seeds with degree ≥ 30 are depicted. Degree was defined as the total number of edges linked to a seed. Seed size is proportional to their degree. Left/right hemisphere is on the left/right, respectively.

Finally, while the consistency analysis revealed some interesting findings, the CPM protocol failed to build models that can predict cumulative standard/extreme bingeing from change in connectivity edges with leave-one-out cross-validation. In particular, model accuracy on test sets: standard bingeing Rpos=-0.03 *(p=* 0.54), Rneg=-0.12 (*p*=0.83); extreme bingeing Rpos=-0.05 (*p*=0.59), Rneg=-0.14 (*p*=0.81).

## 4. Discussion

The current study aimed to examine the longitudinal associations between standard and extreme binge drinking and resting state functional connectivity. When assessing the canonical networks commonly studied in addiction, we found that higher standard bingeing was associated with a decrease in DMN-VAN connectivity from time 1 to time 2, controlling for the initial binge groups at time 1, longitudinal network changes, and other demographic covariates. Although most of the effects of cumulative standard/extreme bingeing did not survive the FDR correction, networks involved in executive function (VAN, CO, Salience, FP) and memory (MTL) were the ones most often associated with binge drinking in this sample. In addition to the network-level analysis, which assumed all edges contribute equally to network connectivity, we also conducted the consistency analysis that focused solely on edges whose change in connectivity strength strongly and consistently associated with bingeing. Although the consistency analysis is exploratory, it allowed for retrieving the edge information that may be lost in the network-level analysis and extended the analysis to non-canonical addiction networks. The consistency analysis identified important seeds with a large number of connectivity edges that showed a consistent pattern of association with bingeing. Indeed, when the whole-brain connectome was considered, both seeds in canonical addiction networks (VAN, DAN, DMN, CO, Salience) and non-canonical networks (Somatomotor Dorsal Network, Auditory) were identified. Lastly, the models from the CPM protocol weren’t able to predict cumulative standard/extreme bingeing from change in connectivity edges in previously unseen data using cross-validation.

Our finding from the network-level analysis was consistent with studies reported the association between binge drinking and abnormal resting state functional connectivity of the DMN (Correas et al., 2016) and the VAN (Herman et al., 2019). Specifically, using MEG to study longitudinal change in resting state connectivity, Correas and colleagues (2016) reported that unlike non-binge drinkers who showed decreased connectivity, college binge drinkers showed increased connectivity in multiple DMN nodes (precuneus, ACC, medial prefrontal cortex, and inferior parietal lobe) across different frequency bands (delta, theta, beta; Cohen’s *d* range 1.4-3.2). Interestingly, this group difference in functional connectivity was not observed for structural connectivity measured with diffusion tensor imaging. The association between binge drinking and DMN connectivity is consistent with what has been reported in the addiction literature (Zhang and Volkow, 2019; Zilverstand et al., 2018). For instance, quantitative metaanalyses have showed that activation in the ACC – a key structure of the DMN – was associated with drug cue onset (Kühn and Gallinat, 2011; Wilson and Sayette, 2015) and self-reported craving (Kühn and Gallinat, 2011). Similarly, resting state connectivity within the DMN has also been reported to correlate with self-reported craving (Li et al., 2016) and degree of relapse (Li et al., 2015). DMN is often thought to have an anti-correlated pattern with networks subserving effortful cognitive processing such as VAN, with the former being task-negative and the latter being task-positive (Raichle, 2015). Researchers have reported that greater binge drinking was associated with decreased resting state connectivity within VAN (Herman et al., 2019). Given the role of VAN in bottom-up, stimulus-driven attentional control (Vossel et al., 2014), Herman and colleagues (2019) postulated that binge drinking could result in attention deficits.

Results from the connectome-based predictive modeling failed to reveal a predictive model (change in connectivity edges predicting cumulative standard/extreme bingeing) that can be generalized to previously unseen data using cross-validation. Several factors may contribute to this finding. Firstly, a recent meta-analysis showed that individual connectivity edge has low test-retest reliability (ICC of 0.29; Noble et al., 2019). Secondly, the strength of univariate association between whole-brain connectome and a behavioral measure of individual differences, in this case standard and extreme bingeing frequency, is often small and might require a larger sample to detect (Marek et al., 2020). Thirdly, although some participants in this sample reported high levels of standard/extreme bingeing, on average, participants had relatively low bingeing frequency (median monthly standard bingeing: sBinge=2.21 days/month, eBinge=2.71 days/month; median monthly extreme bingeing: sBinge=0.33 days/month, eBinge=0.96 days/month), which might explain the weak associations between bingeing and change in connectivity.

The study has several limitations. Firstly, longitudinal standard and extreme bingeing was assessed every 6 months by retrospectively reporting the monthly frequency for the proceeding 6 months, which could make the data subject to memory errors and other self-report biases. Secondly, the binge drinkers in the study were high functioning: the vast majority were still attending college and on average, participants had relatively few bingeing episodes during the two years follow-up. Therefore, different findings might be observed with samples with a heavier or chronic binge drinking pattern.

In summary, our findings suggest that binge drinking is associated with abnormality in networks implicated in allocation of attention control and self-focused processes, which in turn have been implicated in rumination, craving, and relapse. However, the observed effect was relatively weak and was not significantly associated with change in the cognitive performance measures. On one hand, this lack of a connectivity-behavior correlation might reflect our failing to choose and test other, more relevant behavioral correlates of DMN-VAN connectivity for this study. On the other hand, it can also suggest that the effects of binge drinking on neural networks might occur prior to observable behavioral deficits. Future studies are needed to identify cognitive and functional connectivity changes as a function of heavier alcohol binge drinking over longer periods of time.

## CRediT authorship contribution statement

**Tien T. Tong**: Formal analysis, Visualization, Writing – original draft, Writing – review & editing. **Jatin G. Vaidya**: Conceptualization, Funding acquisition, Investigation, Methodology, Data curation, Supervision, Formal analysis, Writing – original draft, Writing – review & editing. **John R. Kramer**: Conceptualization, Investigation, Methodology, Writing – review & editing. **Samuel Kuperman**: Conceptualization, Funding acquisition, Investigation, Methodology, Writing – review & editing. **Douglas R. Langbehn**: Conceptualization, Funding acquisition, Investigation, Methodology, Writing – review & editing. **Daniel S. O’Leary**: Conceptualization, Funding acquisition, Investigation, Methodology, Data curation, Supervision, Project administration, Formal analysis, Writing – original draft, Writing – review & editing.

## Acknowledgements

The work was carried out at University of Iowa Carver College of Medicine, Department of Psychiatry, Iowa City, IA.

## Note

Color should be used for all figures in print

