## SupplementalMaterials for "Impact of Binge Drinking During College on Resting State Functional Connectivity"

**Declarations of interest:** none

**Corresponding Author:**
Tien Tong
200 Hawkins Dr., W278GH, Iowa City, IA 52242

### 1. Descriptions of Neuropsychological Tests

The Stroop task consisted of 4 conditions: 1) read out loud a list of color names printed in black ink, 2) name out loud the colors of a set of colored squares, 3) say out loud the color that a set of non-color words are printed in, and 4) say out loud the color that a set of words are printed in, but the ink color is incongruent with the color word.

In the two in-lab sessions, all subjects passed a breathalyzer test and had negative urine screens for all drugs except for MJ (due to its long half-life)

**3. Details of the Connectome-base Predictive Modeling (CPM) protocol**

For Step 1 (feature selection), edges that were positively or negative correlated with standard/ extreme bingeing (i.e., positive and negative edges) were used in separate models to predict bingeing.

For Step 2 (model fitting), standard/extreme bingeing were the outcome variables, and for each subject, the edge strengths (z-transformed correlation coefficients) were summed across the important edges identified from Step 1 to calculate the predictor of the model.

Model accuracy (Step 3) was the correlation coefficient between the predicted and observed bingeing in the test set across all cross-validation folds (Rpos: coefficient for the model with positive edges, Rneg: coefficient for the model with negative edges).

Since the CPM protocol didn't specify which p-value threshold should be used for Step 1, several thresholds (.01, .001, .0001) were explored. As suggested by Shen and colleagues (2017), $p$<.0001 was chosen, as models using this threshold showed the highest accuracy in predicting bingeing in the training set across all cross-validation folds (see below).

| CPM Models | Mean Rpos Train | Mean Rneg Train |
|---|---|---|
| sBinge: $p$=.01 | 0.71 | 0.69 |
| sBinge: $p$=.001 | 0.73 | 0.71 |
| sBinge: $p$=.0001 | 0.73 | 0.73 |
| eBinge: $p$=.01 | 0.79 | 0.77 |
| eBinge: $p$=.001 | 0.81 | 0.80 |
| eBinge: $p$=.0001 | 0.83 | 0.82 |

The effects of different p-value thresholds on model accuracy were explored using the training sets. sBinge/eBinge: Separate models using change in connectivity edges to predict cumulative standard/extreme bingeing, respectively.

**Supplementary Table 1.** The fixed effect of cumulative bingeing (standard + extreme) on network connectivity.

| | *β* | SE | *t* | DF | Uncorrected *p* | FDR |
|---|---|---|---|---|---|---|
| **MTL** | -0.02 | 0.01 | -2.11 | 182.02 | 0.04 | 0.43 |
| **DMN-VAN** | -0.01 | 0.002 | -3.01 | 172.01 | 0.003 | 0.11 |
| **FP-MTL** | 0.005 | 0.003 | 2.20 | 182.77 | 0.03 | 0.43 |

Network abbreviations: Medial Temporal Lobe (MTL), Default Mode Network (DMN), Ventral Attention Network (VAN), and Fronto-Parietal (FP).

**Supplementary Table 2.** Spearman's correlation coefficients between change (d = Time 2 minus Time 1) in cognitive performance and cumulative standard and extreme bingeing.

| | d Stroop Effect 1 | d Stroop Effect 2 | d Digit Span Total Correct | d Letter Fluency Total Correct | d Trail Making Elapsed Time (B-A) |
|---|---|---|---|---|---|
| Cumulative sBinge | -0.14 | -0.09 | 0.03 | -0.02 | 0.04 |
| Cumulative eBinge | -0.14 | -0.08 | 0.06 | 0.02 | 0.12 |

sBinge: log-transformed cumulative standard bingeing; eBinge: log-transformed cumulative extreme bingeing. No correlation was significant.

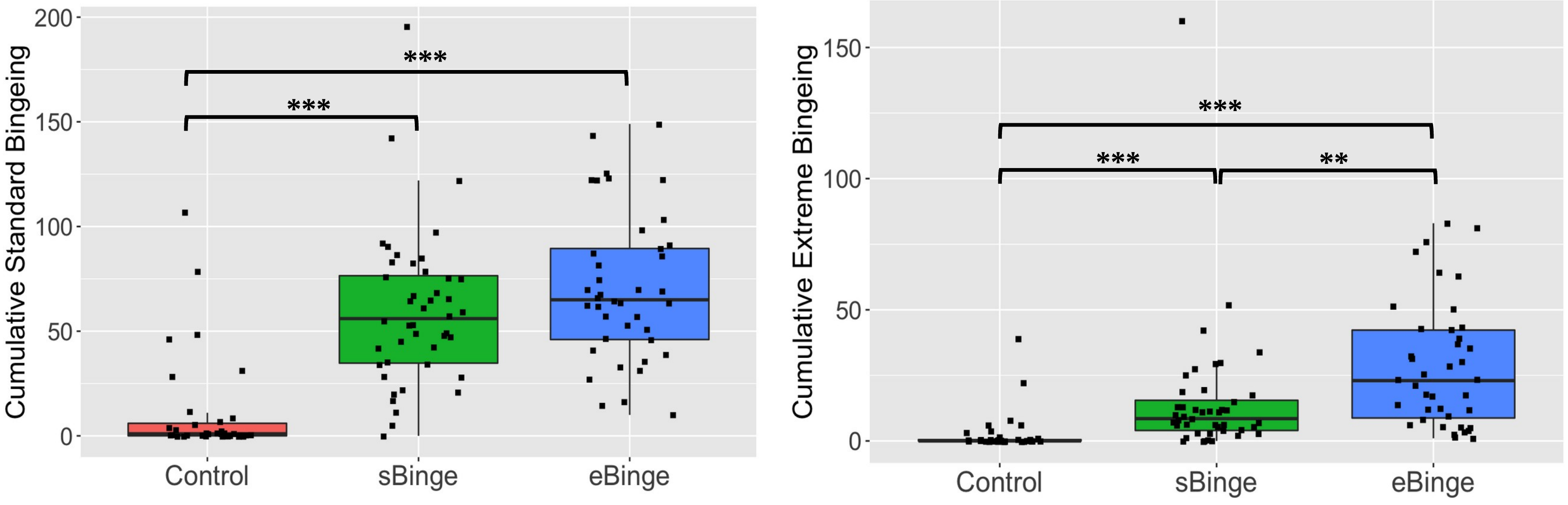

Cumulative Standard Bingeing
200
150
100
50
0
***
***
Control
sBinge
eBinge
Cumulative Extreme Bingeing
150
100
50
0
***
***
**
Control
sBinge
eBinge
